# A normative account of confirmation bias during reinforcement learning

**DOI:** 10.1101/2020.05.12.090134

**Authors:** Germain Lefebvre, Christopher Summerfield, Rafal Bogacz

## Abstract

Reinforcement learning involves updating estimates of the value of states and actions on the basis of experience. Previous work has shown that in humans, reinforcement learning exhibits a confirmatory bias: when updating the value of a chosen option, estimates are revised more radically following positive than negative reward prediction errors, but the converse is observed when updating the unchosen option value estimate. Here, we simulate performance on a multi-arm bandit task to examine the consequences of a confirmatory bias for reward harvesting. We report a paradoxical finding: that confirmatory biases allow the agent to maximise reward relative to an unbiased updating rule. This principle holds over a wide range of experimental settings and is most influential when decisions are corrupted by noise. We show that this occurs because on average, confirmatory biases lead to overestimating the value of more valuable bandits, and underestimating the value of less valuable bandits, rendering decisions overall more robust in the face of noise. Our results show how apparently suboptimal learning policies can in fact be reward-maximising if decisions are made with finite computational precision.

## Introduction

Confirmation bias refers to seeking or interpreting evidence in ways that are influenced by existing beliefs, and it is a ubiquitous feature of human perceptual, cognitive and social processes, and a longstanding topic of study in psychology (Nickerson, 1998). Confirmatory biases can be pernicious in applied settings, for example when clinicians overlook the correct diagnosis after forming a strong initial impression of a patient (Groopman, 2007). In laboratory, confirmation bias has been studied with a variety of paradigms (Nickerson, 1998; Talluri, Urai, Tsetsos, Usher, & Donner, 2018). One paradigm in which the confirmation bias can be observed and measured involves reinforcement learning tasks, where participants have to learn from positive or negative feedback which options are worth taking (Chambon et al., 2019; Palminteri, Lefebvre, Kilford, & Blakemore, 2017), and this paper focusses on confirmation bias during reinforcement learning.

In the laboratory, reinforcement learning is often studied via a “multi-armed bandit” task in which participants choose between two or more states that pay out a reward with unknown probability (Daw, O’Doherty, Dayan, Seymour, & Dolan, 2006). Reinforcement learning on this task can be modelled with a simple principle known as a delta rule (Rescorla & Wagner, 1972), in which the estimated value 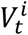 of the chosen bandit ***i*** on trial ***t*** is updated according to:

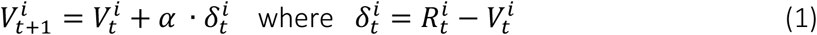

In the above equation, 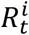 is the payout for option ***i*** on trial ***t***, 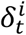 is the reward prediction error, and ***α*** is a learning rate in unity range. If ***α*** is sufficiently small, 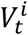 tend to converge over time to the vicinity of expected value of bandit *i* (in stationary environments).

This task and modelling framework have also been used to study the biases that humans exhibit during learning. One line of research has suggested that humans may learn differently from positive and negative outcomes. For example, variants of the model above which include distinct learning rates for positive and negative updates to 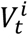 have been observed to fit human data from a 2-armed bandit task better, even after penalising for additional complexity (Gershman, 2015; Niv, Edlund, Dayan, & O’Doherty, 2012). When payout is observed only for the option that was chosen, updates seem to be larger when the participant is positively rather than negatively surprised, which might be interpreted as a form of optimistic learning (Lefebvre, Lebreton, Meyniel, Bourgeois-Gironde, & Palminteri, 2017). However, a different pattern of data was observed in follow-up studies in which counterfactual feedback was also offered – i.e., the participants were able to view the payout associated with both chosen and unchosen options. Following a feedback on the unchosen option, larger updates were observed for negative prediction errors (Chambon et al., 2019; Palminteri et al., 2017; Schuller et al., 2020). This is consistent with a confirmatory bias rather than a strictly optimistic bias, whereby belief revision helps to strengthen rather than weaken existing preconceptions about which option may be better.

One obvious question is why confirmatory biases persist as a feature of our cognitive landscape – if they promote suboptimal choices, why have they not been selected away by evolution? One variant of the confirmation bias, a tendency to overtly sample information from the environment that is consistent with existing beliefs, has been argued to promote optimal data selection: where the agent chooses its own information acquisition policy, exhaustively ruling out explanations (however obscure) for an observation would be highly inefficient (Oaksford & Chater, 2003). However, this account is unsuited to explaining the differential updates to chosen and unchosen options in a bandit task with counterfactual feedback, because in this case feedback for both options is freely displayed to the participant, and there is no overt data selection problem.

It has been demonstrated previously that biased estimates of value can paradoxically be beneficial in two-armed tasks in the sense that under standard assumptions, they maximise the average total reward for the agent (Caze & van der Meer, 2013). This happens because with such biased value estimates, the difference 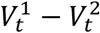 may be magnified, so with a noisy choice rule (typically used in reinforcement learning models) the option with the higher reward probability is more likely to be selected. Caze and van der Meer (2013) considered a standard reinforcement learning task in which feedback is provided only for the chosen option. In that task the reward probabilities of the two options in the task determine whether it is beneficial to have higher learning rate after positive or negative prediction error (Caze & van der Meer, 2013). In other words, when only outcome of chosen option is observed, optimistic bias is beneficial for some reward probabilities, while pessimistic bias for other.

In this paper we show that if the participants are able to view the payouts associated with both chosen and unchosen options, reward is typically maximized if the learning rates follow the pattern of the confirmation bias, i.e. are higher when the chosen option is rewarded and unchosen option is unrewarded. We find that this benefit holds over a wide range of settings, including both stationary and nonstationary bandits, with different reward probabilities, across different epoch lengths, and under different levels of choice variability. We also demonstrate that such confirmation bias tends to magnify the difference 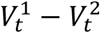 and hence makes the choice more robust to the decision noise. These findings may explain why the humans tend to revise beliefs to a smaller extent when outcomes do not match with their expectations.

Below we formalize the confirmation bias in a reinforcement learning model, compare its performance in simulations with models without confirmation bias, and formally characterise the biases introduced in value estimates. We also point that the confirmation bias not only typically increases the average reward, but may increase the reward rate to even higher extent.

## Reinforcement learning models

### Confirmation model

We analyse properties of a *confirmation* model (Palminteri et al., 2017), which describes learning in a two-armed bandit task where feedback is provided for both options on each trial. The model updates the corresponding value estimates 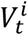 according to a delta rule with two learning rates: ***α^C^*** for confirmatory updates (i.e. following positive prediction errors for the chosen option, and negative for the unchosen option) and ***α^D^*** for disconfirmatory updates (i.e. following negative prediction errors for the chosen option, and positive for the unchosen option) (Palminteri et al., 2017). Thus on each trial ***t***, if the agent chooses the option 1, the model updates the values 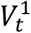 and 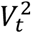 of the chosen and unchosen options respectively, such that:

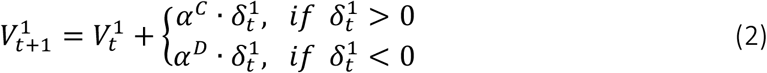

and

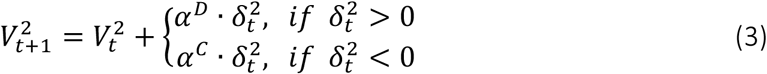

with 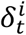 being the prediction error for bandit ***i*** on trial ***t*** defined in Equation 1. We define an agent with a confirmatory bias as one for whom ***α^C^ > α^D^***, whereas an agent with a disconfirmatory bias has ***α^C^ < α^D^***, and an agent with no bias (or a neutral setting) has ***α^C^ = a^D^***. Note that for ***α^C^* = *α^D^* = *α***, the model amounts to a standard delta-rule model with a unique learning rate *α* defined in Equation 1, and we refer to it as *unbiased*.

### Decaying learning rate model

We compared the performance of the *confirmation* model in a stable environment to an optimal value estimator which for each option computes the average of rewards seen so far. Such values can be learned by a model using the update given in Equation 1 with the learning rate ***α*** decreasing over trials according to 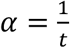, where ***t*** is the trial number (note that with the counterfactual feedback, *t* is also equal to the number of times the reward for this option has been observed).

### Decision Policies

In this paper we consider three policies for making a choice on the basis of learned values: *hardmax, softmax* and *ε-greedy* policies. The hardmax is a noiseless policy selecting deterministically the arm associated to the highest value. The softmax is a probabilistic action selection process associating to each arm *a* the probability 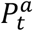 of being selected based on their respective values such that:

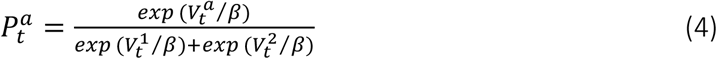

In the above equation ***β*** is the temperature of the *softmax* function, and the higher the temperature, the more random the decision is. To mathematically analyse properties of the confirmation model, we also consider a simpler stochastic choice rule ε-greedy, which on majority of trials selects option with highest estimated value, while with certain fixed probability selects an action randomly.

## Effects of confirmation bias on average reward

### Methods of simulation

Our goal was to test how outcomes vary with a confirmatory, disconfirmatory or neutral bias across a wide range of different settings that have been the subject of previous empirical investigation in humans and other animals. We considered the tasks involving choice between two options. Each bandit ***i*** may yield reward ***R* = 1** with probability ***p^i^***, and no reward (***R* = 0**) with probability **1 – *p^i^***. Importantly we assumed that the agent observes on each trial the payouts for both options: the chosen one and the not chosen option (**Fig. 1a**). We consider an agent who chooses among bandits for **2^*n*^** trials, where ***n*** varied from 2 to 10 in simulations (**Fig. 1b**), and the presented rewards were averaged over these values of ***n*** (unless otherwise stated).

**Figure 1.**
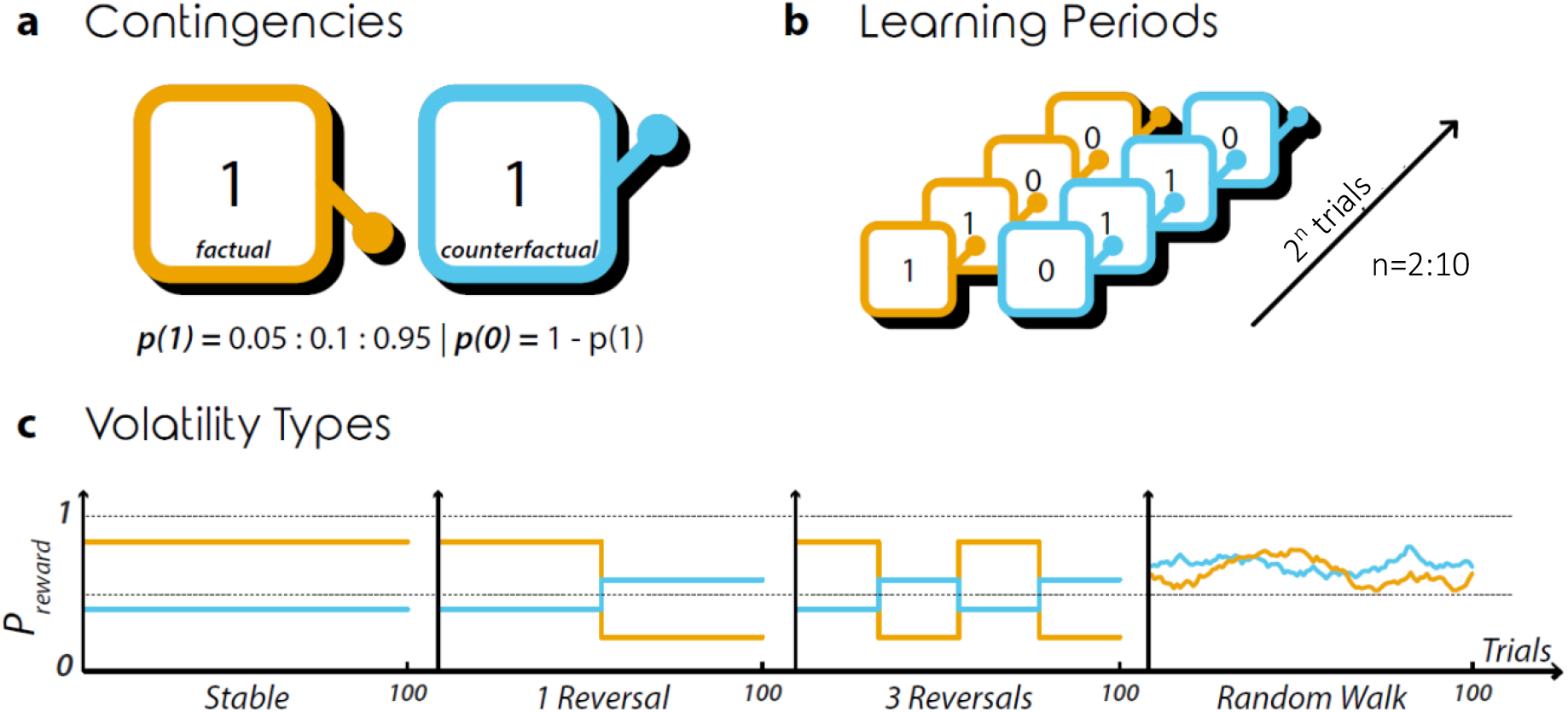
Simulation Setup. a. Reward contingencies. The illustration represents the chosen (orange) and unchosen (blue) bandits each with a feedback signal (central number). Below, we state the range of possible outcomes and probabilities. **b**. Learning Periods. The illustration represents the different length of the learning period and the different outcomes combinations potentially received by the agents. **c**. Volatility Types. The line plots represent the evolution of the two arms probability across trials in the different volatility conditions.

We considered four different ways in which the reward probabilities ***p^i^*** are set, illustrated schematically in **Fig. 1c**. First we considered *stable* environments in which reward probabilities were constant. We also considered *1 reversal* and *3 reversals* conditions where the payout probabilities were reversed to **1 – *p^i^*** once in the middle of the task (second display in **Fig. 1c**), or three times at equal intervals (third display in **Fig. 1c**). In *stable, 1 reversal* and *3 reversals* conditions, the initial probabilities ***p^i^*** at the start of the task were sampled at intervals of 0.1 in the range [0.05, 0.95] such that ***p*^1^ ≠ *p*^2^**, and we tested all possible combinations of these probabilities (that is **45** probability pairs). Unless otherwise noted, results are averaged across these initial probabilities.

Additionally, we considered *random walk* condition where the probabilities were initialized to a random number from uniform distribution on interval from 0 to 1, and then drifted over trials as follows:

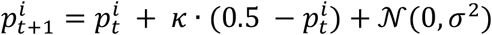

with ***κ*** being a parameter decaying the reward probability towards **0.5** (here set to ***κ* = 0.001**) and ***σ*** being the standard deviation of the normal distribution from which the fluctuations in probabilities were sampled (here set to ***σ* = 0.02**). Sample probabilities generated from this process are shown in the fourth display in **Fig. 1c**.

We conduct all simulations numerically, sampling the initial payout probabilities and experiment length(s) exhaustively, varying ***α^C^*** and ***α^D^*** exhaustively, and noting the average reward obtained by the agent in each setting. The model is simulated with all possible combinations of learning rates ***α^C^*** and ***α^D^*** defined in the range **[0.05,0.95]** with increments of **0.1**, that is **19^2^** learning rate combinations. For each combinations of parameters, the simulations were performed 1000 times for all but the random walk condition where simulations are performed 100000 times to account for the increased variability. Results are averaged for plotting and analysis. In all cases, inferential statistics were conducted using nonparametric tests with an alpha of p < 0.001 and Bonferroni correction for multiple comparisons.

### Results of simulations

**Fig. 2** plots total reward obtained in the stationary bandit problem as a function of ***α^C^*** (y-axis) and ***α^D^*** (x-axis), for the sequence length of 1024 and averaged across payout probabilities, for both the hardmax (left) and softmax (right) rules. The key result is that rewards are on average greater when ***α^C^ > α^D^*** (warmer colours above the diagonal) relative to when they are equal or when ***α^C^ < α^D^***. We tested this finding statistically by repeating our simulations multiple times with resampled stimulus sequences (and choices in the softmax condition) and comparing the accrued reward to a baseline in which ***α^C^* = *α^D^* = 0.05**, i.e. the most promising unbiased setting for ***α***. The area enclosed by black line in **Fig. 2a-b** indicate combinations of learning rates that yield rewards higher than the unbiased setting. **Fig. 2b** confirms that in particular for the more plausible case where decisions are noisy (i.e. softmax temperature ***β* > 0**), there is a reliable advantage for a confirmatory update policy in the bandit task.

**Figure 2.**
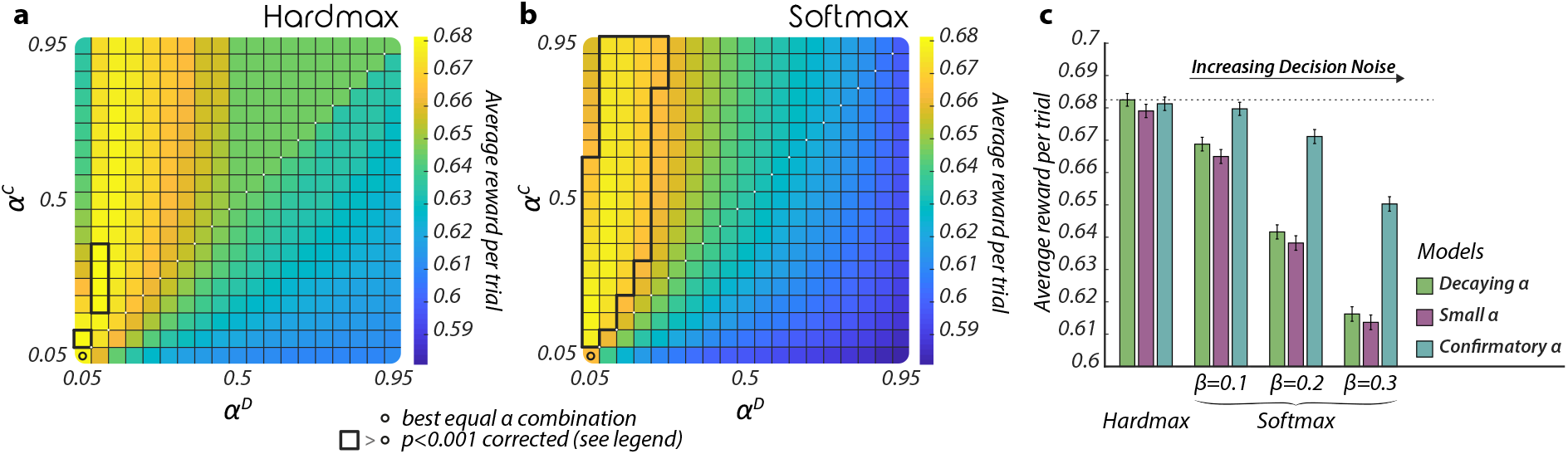
Dependence of reward on learning rate and decision noise in a stable environment. **a** and **b**. Average reward for all learning rate combinations. The heatmaps represent the per trial average reward for combinations of ***α^C^*** (y-axis) and ***α^D^*** (x-axis), averaged across all reward contingencies and agents in the stable condition with 1024 trials. Areas enclosed by black lines represent learning rate combinations for which the reward is significantly higher than the performance of the best equal learning rates combination represented by a black circle, one-tailed independent samples rank-sum tests, p<0.001 corrected for multiple comparison. **a**. Deterministic Decisions. Simulated reward is obtained using a noiseless *hardmax* policy. **b**. Noisy Decisions. Simulated reward is obtained using a noisy *softmax* policy with ***β* = 0.1**. **c**. Comparison with optimal models. The bar plot represents the per trial average reward of the *confirmation* model, the *small learning rate* model and the *decaying learning rate* model for four different levels of noise in the decision process. In simulations of the *confirmation model*, the best learning rates combination was used for each noise level (i.e. ***α^C^* = [0.1,0.15,0.3,0.35]** and ***α^D^* = 0.05**). Bars represent the means and error bars the standard deviations across agents, all reward levels are significantly different from each other, two-tailed independent samples rank-sum tests, p<0.001.

We compared the performance of the confirmation model to the decaying learning rate model described above, which maximizes reward under the assumption that payout probabilities are stationary and decisions are noiseless (i.e. under a hardmax choice rule). We confirmed this by plotting the average reward under various temperature values for three models: one in which a single learning rate was set to a fixed low value ***α* = 0.05** (*small learning rate* model) one in which it was optimally annealed (*decaying learning rate* model), and one in which there was a confirmatory bias (*confirmation* model; **Fig. 2c**). As can be seen, only under ***β* = 0** the confirmation bias does not increase rewards; as soon as decision noise increases, the relative merit of the confirmation model grows sharply. Importantly, whereas the performance advantage for the decaying learning rate model in the absence of noise (under ***β* = 0**) was very small (on the order of 0.2%), the converse advantage for the confirmatory bias given noisy decisions was numerically larger (1.6%, 4.6% and 5.5% under ***β* = 0.1,0.2, 0.3** respectively).

Next, we verified that these results held over different trial lengths and for differing volatility conditions. The results (averaged over different numbers of trials) are shown in **Fig. 3**. One can see equivalent results presented for a paradigm involving stable contingencies (**Fig. 3a and 3e**), a reversal of probability between the two bandits midway through the sequence (**Fig. 3b and 3f**), for three such reversals (**Fig. 3c and 3g**), and for a random walk in which probabilities drift upwards or downwards on each trial (**Fig. 3d and 3h**). When decisions are noisy, in all four cases, confirmatory agents reap more rewards than disconfirmatory agents, and also than agents for whom there is a single ***α*** selected to maximise reward (**Fig. 3e-h**). When the decisions are based on the hardmax choice rule, there was no biased combination of learning rates giving significantly higher rewards than unbiased model (**Fig. 3a-d**). Nevertheless, there still existed combinations of parameters with ***α^C^* > *α^D^*** yielding reward similar to that from the unbiased model.

**Figure 3.**
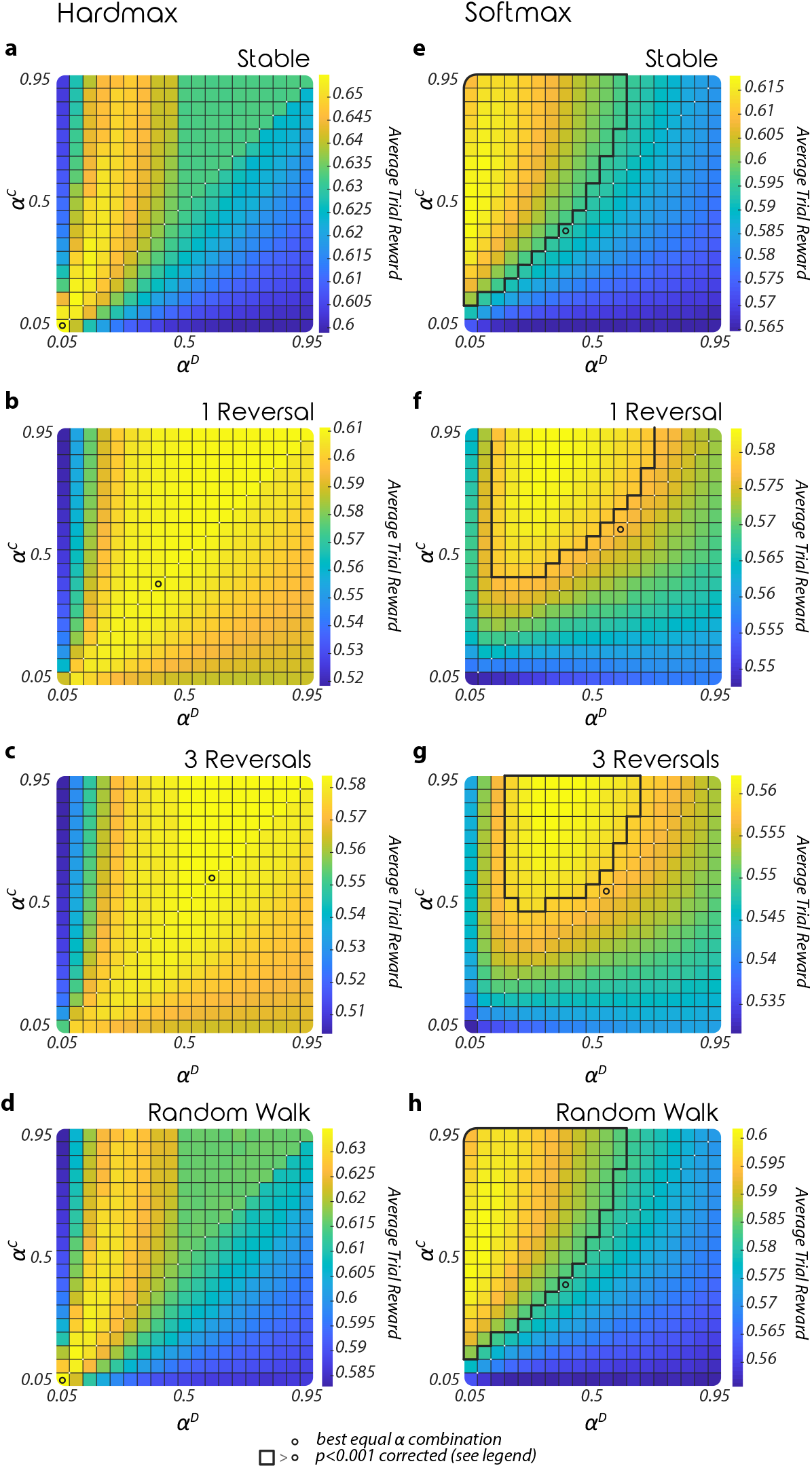
Dependence of reward on learning rate and decision noise in different environments. **a, b, c, d, e, f, g & h**. The heatmaps represent the per trial average reward for combinations of ***α^C^*** (y-axis) and ***α^D^*** (x-axis) given a *hardmax* policy (**a, b, c, d**) or a *softmax* policy (***β* = 0.3**) (**e, f, g, h**). The performance is averaged across all reward contingencies, period lengths and 1000 agents in the stable condition (**a, e**), 1 reversal condition (**b, f**), 3 reversals condition (**c, g**) or 100000 agents in the random walk condition (**d, h**). Areas enclosed by black lines represent learning rate combinations for which the reward is significantly higher than the reward of the best equal learning rates combination represented by a black circle, one-tailed independent samples rank-sum tests, ***p* < 0.001** corrected for multiple comparisons.

Subsequently, we tested how the sequence length affected the relative advantage conferred by a confirmatory bias. In **Fig. 4a**, we show that the advantage for the confirmatory over the unbiased model holds true for all but the very shortest sequences and continues to grow up to sequences of 1024 trials. Finally, the confirmatory model is most advantageous at intermediate levels of decisions noise (as quantified here by the softmax temperature). As we have seen, the relative numerical and statistical advantage is lower if we assume no decision noise, but as decision noise grows to the extent that performance tends towards random, all differences between different update policies disappear (**Fig. 4b**).

**Figure 4.**
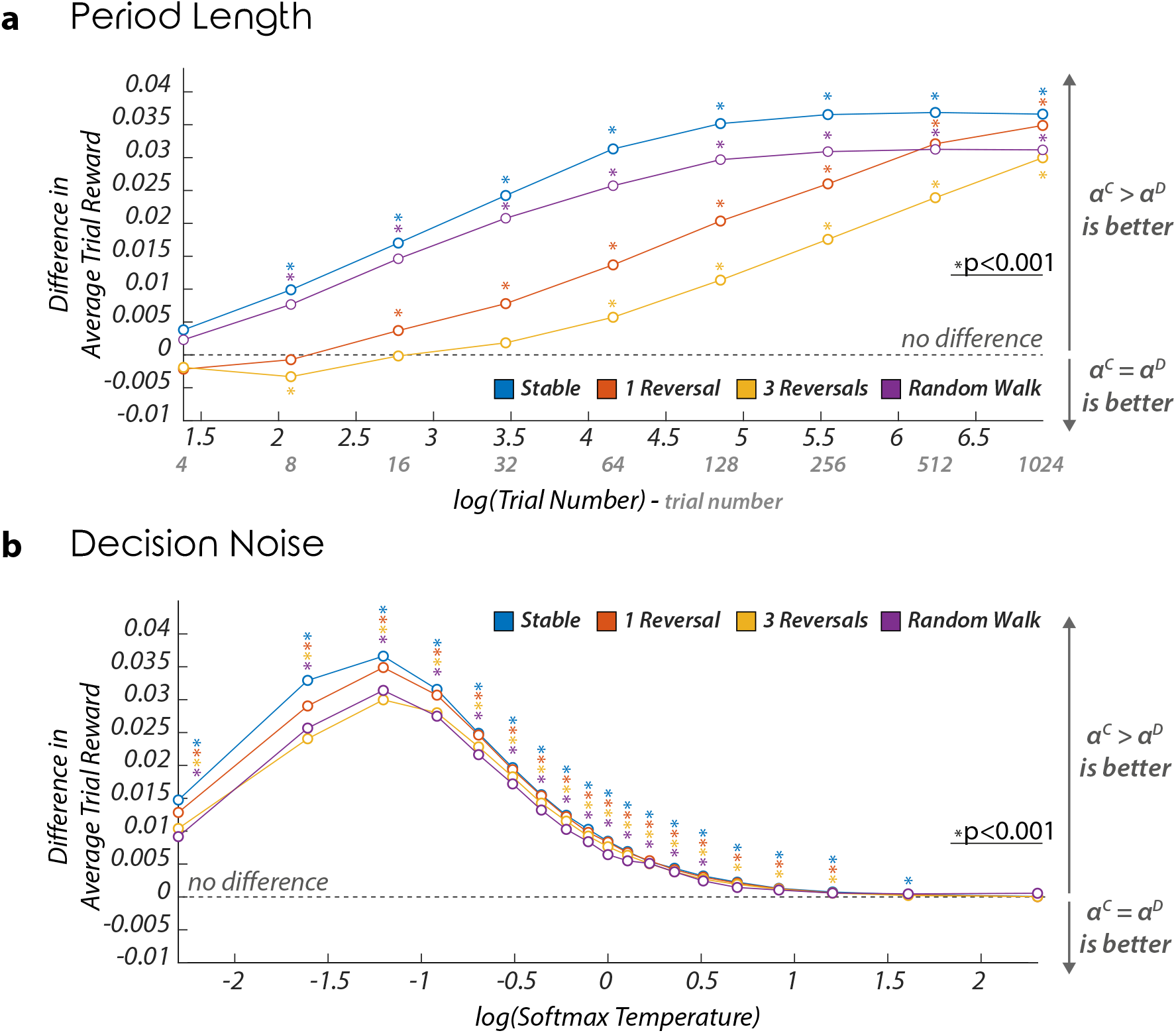
Effects of period length and decision noise on the relative performance of confirmation model. **a**. Effect of period length on reward. The line plot represents the difference in average reward between the confirmation model (with the best confirmatory learning rate combination per period) and the unbiased model (with the best per period single learning rate) in function of the log of the period length, and for the four different volatility conditions. The logarithmic transformation of the trial number is for illustrative purpose only. *, ***p* < 0.001**, two-tailed independent rank-sum tests. **b**. Effect of decision noise on performance. The line plot represents the difference in per trial average performances of the confirmation model (with the best confirmatory learning rates combination) and the unbiased model (with the single best learning rate) in function of the log of *softmax* temperature, and for the four different volatility conditions. The logarithmic transformation of the *softmax* temperature is for illustrative purpose only. *, ***p* < 0.001**, two-tailed independent rank-sum tests.

**Fig. 5a-c** show that confirmation bias increases average reward irrespectively of the range of reward probabilities for the two options. The consistent effect of confirmation bias contrasts with the effects of biases in learning rates found in a previous study mentioned in the Introduction (Caze & van der Meer, 2013), and we will come back to comparison of these studies in the Discussion. **Fig. 5d-f** show that confirmation bias also increases the average outcome in the aversive tasks in which the outcome on each trial 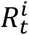 is either −1 or 0.

**Figure 5.**
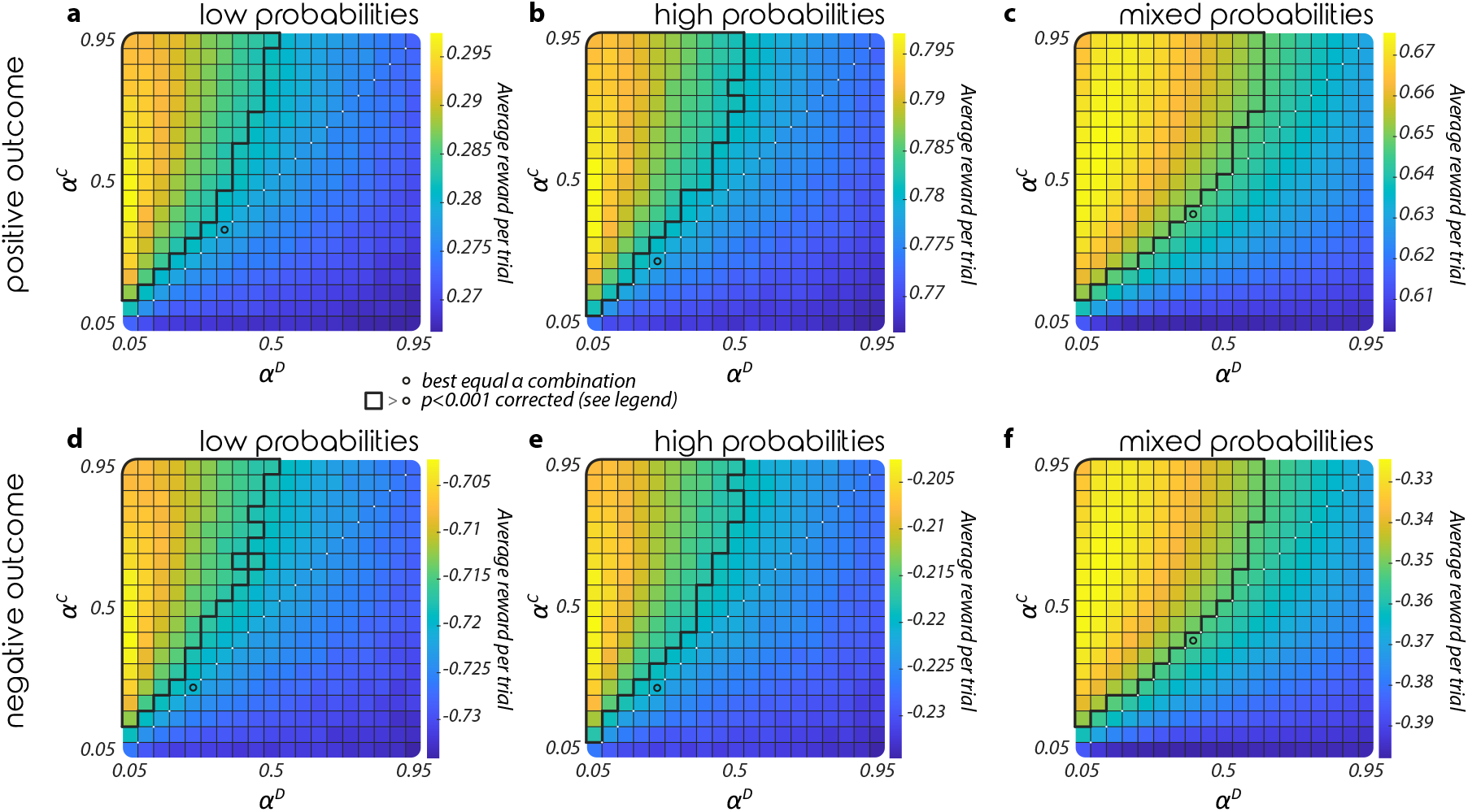
Dependence of reward on learning rate, outcome valence and reward contingencies. **a, b, c, d, e, & f**. The heatmaps represent the per trial average reward for combinations of ***α^C^*** (y-axis) and ***α^D^*** (x-axis) given a *softmax* policy (***β* = 0.3**) in a stable environment. The performance is averaged across 1000 agents, all period lengths and low reward contingencies, i.e. ***p*^1^ < 0.5** and ***p*^2^ < 0.5** (**a, d**), high reward contingencies, i.e. ***p*^1^ > 0.5** and ***p*^2^ > 0.5** (**b, e**), mixed reward contingencies, i.e. ***p*^1^ > 0.5** and ***p*^1^ < 0.5** (**c, f**). Outcome valence was either positive, i.e. [0 1] (**a, b, c**) or negative, i.e. [-1 0] (**d, e, f**). Areas enclosed by black lines represent learning rate combinations for which the reward is significantly higher than the reward of the best equal learning rates combination represented by a black circle, one-tailed independent samples rank-sum tests, ***p* < 0.001** corrected for multiple comparisons.

## Confirmation bias magnifies difference between estimated values

The above simulations show that a confirmatory update strategy – one which privileges the chosen over the unchosen option – is reward-maximising across a wide range of experimental conditions, in particular when decisions are noisy. Why would this be the case? It is well known, for example, that adopting a single small value for ***α*** will allow value estimates to converge to their ground truth counterparts. Why would an agent want to learn biased value estimates? To answer this question, we demonstrate below that the confirmation bias often magnifies the differences between estimated values and hence make choices more robust to decision noise. We first show it on an intuitive example and then more formally.

### Example of the effects of confirmation bias

We selected three parametrisations of the update rules and examined their consequences in more detail. The selected pairs of values for ***α^C^*** and ***α^D^*** are illustrated in **Fig. 6a** (symbols Δ, × and o). The first corresponded to an unbiased update rule: ***α^C^* = *α^D^* = 0.25**; the second to a moderately biased rule (***α^C^* = 0.35, *α^D^* = 0.15**); and the third to a severely biased rule (***α^C^* = 0.45, *α^D^* = 0.05**). Let us refer to the bandit with a higher reward probability as richer and to the other bandit as poorer. We chose a setting in which reward probability for the richer bandit is ***p*^+^ = 0.65**, while for the poorer bandit it is ***p*^−^ = 0.35**.

**Figure 6.**
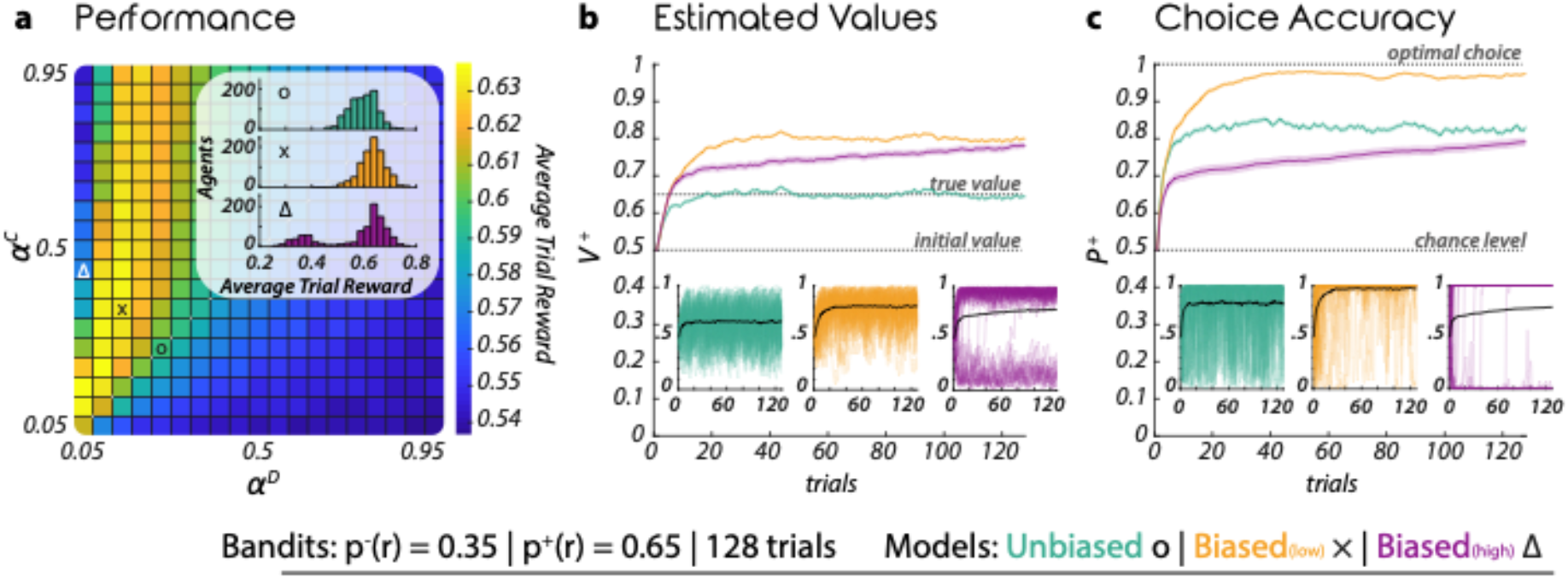
Mechanism by which confirmation bias tends to increase reward. **a**. Average reward and reward distributions for different levels of confirmation bias. The heatmap represents the per trial average reward of the confirmation model for all learning rates combinations (confirmatory learning rates are represented on the y-axis whereas disconfirmatory learning rates are represented on the x-axis) associated with a *softmax* policy with ***β* = 0.1**. The rewards concern the stable condition with 128 trials and asymmetric contingencies (***p*^-^ = 0.35** and ***p*^+^ = 0.65**) and are averaged across agents. The three signs inside the heatmap (**Δ**, x and +) represent the three learning rates combinations used in the simulations illustrated in panels **b** and **c**. The histograms show the distribution across agents of the average per trial reward for the three different combinations. **b**. Estimated values. The line plots represent the evolution of the best option value ***V*^+^** across trials. The large plot represents the agents-averaged value of the best option across trials for three different learning rates combinations, “unbiased” (***α^C^* = *α^D^* = 0.25**), “biased (low)” (***α^C^* = 0.35** and ***α^D^* = 0.15**) and “biased (high)” (***α^C^* = 0.45** and *α^D^* = 0.05). The lines represent the mean and the shaded areas, the SEM. The small plots represent the value of the best option across trials plotted separately for the three combinations. The thick lines represent the average across agents and the lighter lines the individual values of 5% of the agents. **c**. Choice Accuracy. The line plots represent the evolution of the probability to select the best option across trials. The large plot represents the agents-averaged probability to select the best option across trials for three different learning rates combinations, “unbiased” (***α^C^* = *α^D^* = 0.25**), “biased (low)” (***α^C^* = 0.35** and ***α^D^* = 0.15**) and “biased (high)” (***α^C^* = 0.45** and ***α^D^* = 0.05**). The lines represent the mean and the shaded areas, the SEM. The small plots represent the probability to select the best option across trials plotted separately for the three combinations. The thick lines represent the average across agents and the lighter lines the individual probability for 5% of the agents.

For each update rule, we plotted the evolution of the value estimate for the richer bandit ***V*^+^** over trials (**Fig. 6b**) as well as aggregate choice accuracy (**Fig. 6c**). Beginning with the choice accuracy data, one can see that intermediate levels of bias are reward-maximising, in the sense that they increase the probability that the agent chooses the bandit with the higher payout probability, relative to an unbiased or a severely biased update rule (**Fig. 6c**). This is of course simply a restatement of the finding that biased policies maximise reward (see shading in **Fig. 6a**). However, perhaps more informative are the value estimates for ***V*^+^** under each update rule (**Fig. 6b**). As expected, the unbiased learning rule allows the agent to accurately learn the appropriate value estimate, such that after a few tens of trials, ***V*^+^ ≈ *p*^+^ = 0.65** (grey line). By contrast, the confirmatory model *overestimates* the value of the richer option (converging close to ***V*^+^~0.8** despite ***p*^+^ = 0.65**, and (not shown) the model *underestimates* the value of the poorer option ***p*^−^ = 0.35**). Thus, the confirmation model outperforms the unbiased model despite misestimating the value of both the better and the worse option. How is this possible?

To understand this phenomenon, it is useful to consider the policy by which simulated choices are made. In the two-armed bandit case, the softmax choice rule of Equation 4 can be re-arranged to the following logistic function:

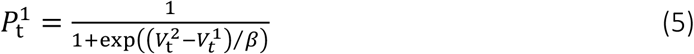

Here, the choice probability depends both on the inverse slope of the choice function ***β*** and the difference in value estimates for bandits 1 and 2. The effect of the confirmation bias is to inflate the quantity 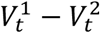 away from zero in either the positive or the negative direction, thereby ensuring choice probabilities that are closer to 0 or 1 even in the presence of decision noise (i.e. larger ***β***). This comes at a potential cost of overestimating the value of the poorer option rather than the richer, which would obviously hurt performance. The relative merits of an unbiased vs. biased update rule are thus shaped by the relative influence of these factors. When the rule is unbiased, the model does not benefit from the robustness conferred by inflated value estimates. When the model is severely biased, the probability of confirming the incorrect belief is excessive – leading to a high probability that the poorer option will be overvalued rather than the richer (see the bimodal distribution of value estimates in **Fig. 6b**, inset). Our simulations show that when this happens, the average reward is low, resulting in bimodal distribution of rewards across simulations (inset in **Fig. 6a**). However, there exists a “goldilocks zone” for confirmatory bias in which the benefit of the former factor outweighs the cost of the latter. This is why a confirmation bias can help maximise reward.

### Analysis of biases in estimated values

This section shows formally that under certain assumptions the confirmation bias increases the distance between the estimated values. We followed the approach from a previous study analysing biases in values due to unequal learning rates (Caze & van der Meer, 2013) and analysed the values learned in a stable environment. Due to stochastic nature of rewards in the task, 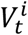 constantly fluctuate, but with time they approach a vicinity of values known as a stochastic fixed points 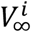, in which they will not change on average, i.e. 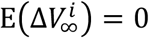.

We were not able to obtain tractable analytic expressions for stochastic fixed points of values when the softmax choice rule was assumed, hence we considered a simpler ε-greedy choice rule. We denote the probability of selecting an option with a lower estimated value by *ε*. For simplicity, let us assume that a learned value for the richer option 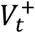 is higher than the value for the poorer option 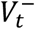. This assumption tends to be correct when ***p*^+^** is sufficiently higher ***p*^−^**, but it may not be satisfied when ***p*^+^** and ***p*^−^** are similar, because of fluctuations in values occurring during learning and a possibility that the model is stuck in preferring poorer option due to the confirmation bias. Under this assumption the richer option is selected with probability **1 – *ε***, while the poorer option with probability *ε*.

The average change in the value of the richer option is then given by:

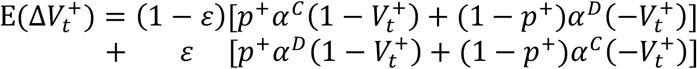

In the above equation the first line corresponds to changes occurring when the richer option is chosen, and the second line when the poorer option is chosen. Within each line, the first term in a square bracket corresponds to a change when the richer option yields rewards, while the second term when the richer option is not rewarded. To find the value in a stochastic fixed point, we set the left hand side of the above equation to 0 (because the stochastic fixed point is defined as the value in which the average value change is 0), and solve for 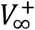 to obtain:

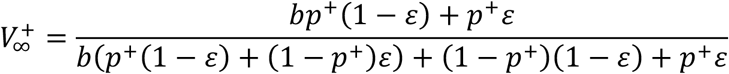

where 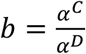 quantifies the amount of confirmation bias. The above equation shows that the value 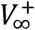 in a stochastic fixed point does not depend on the individual learning rates, but only on their ratio, similarly as in a previous study (Caze & van der Meer, 2013). Analogous analysis shows that the stochastic fixed point for the poorer option is equal to:

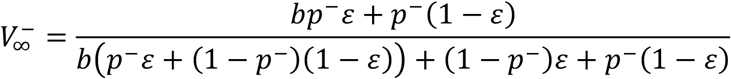

We now demonstrate that the confirmation bias increases 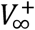 and decreases 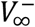. To evaluate the effect to increasing ***α^C^*** relatively to ***α^D^*** on 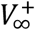, we compute

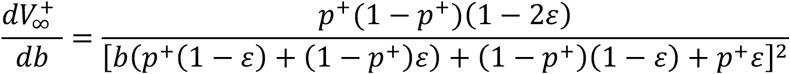

The above expression is non-negative because the denominator is non-negative (as it is a square) and the numerator is a product of non-negative terms. This derivative will be positive if **0 < *p*^+^ < 1**, and 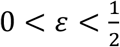, i.e. when the rewards are non-deterministic, and the choice policy is stochastic but not completely random. The derivative for 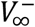 is equal to an analogous expression but with a negative sign:

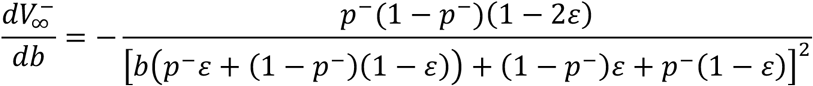

In summary, for stochastic rewards and stochastic choice policy, the confirmation bias increases 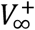 and decreases 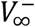, and hence it magnifies the difference between the values.

**Fig. 7** compares theoretically predicted difference between the learned values (dashed lines) with the results of simulations. The simulations match experimental predictions well except for when ***p*^+^** and ***p*^−^** are equal or close to each other (in **Fig. 7a** and **7c**), because then the assumption that ***V*^+^ > *V*^−^** stated at the start of the section is no longer always valid. For the unbiased model (green curves), the difference in estimated values follow the difference in reward probabilities. Importantly, when ***p*^+^** and ***p*^−^** differ, the confirmation bias increases the difference between the learned values (except for the case where both outcomes are deterministic, i.e. ***p*^+^ = 1** and ***p*^−^ = 0** in **Fig. 7a**).

**Figure 7.**
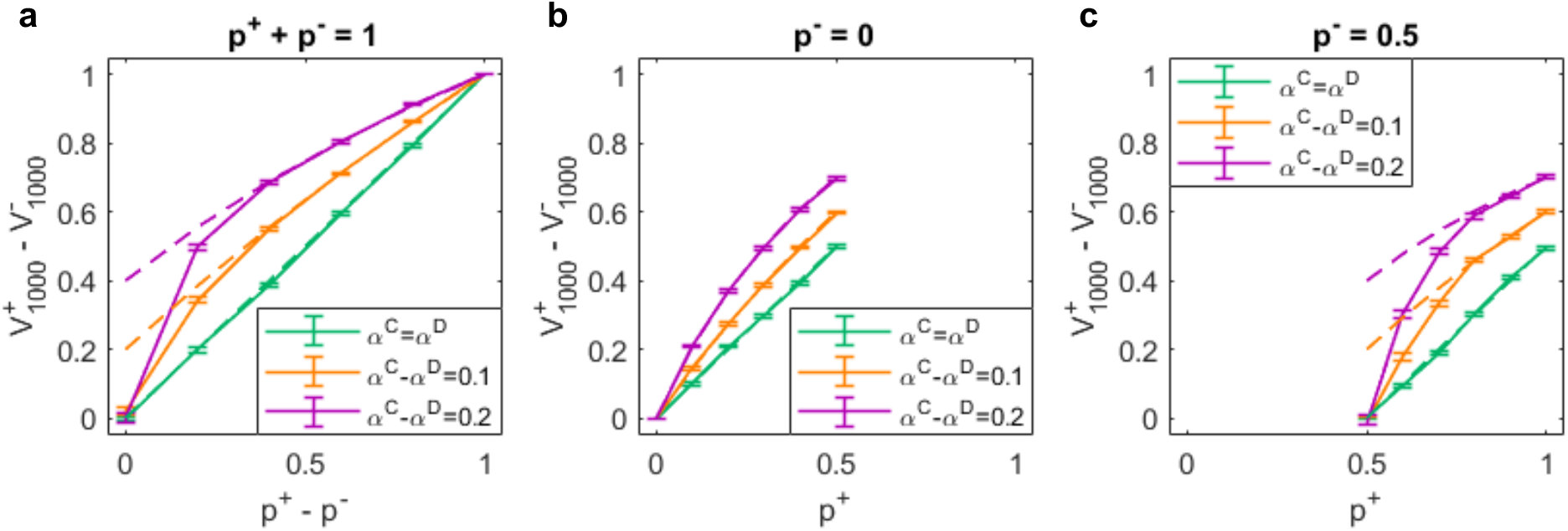
Differences between learned values of the richer and poorer options. Dashed lines show theoretically predicted difference. Solid lines were obtained in simulations, and show the difference between the learned values after 1000 trials in a stable environment. These differences are averaged over 1000 simulations and error bars show standard error. Different colours correspond to different combinations on learning rates (shown in key). X-axes show the difference in reward probabilities of the richer and poorer options in panel a, and the reward probability for the richer option in the other panels. Different panels correspond to different constraints on reward probabilities, stated above each panel.

## Effects of confirmation bias on reward rate

The analysis shown in **Fig. 6** illustrates why the benefit confirmation drops off as the bias tends to the extreme – it is because under extreme bias, the agent falls into a feedback loop whereby it confirms its false belief that the lower-valued bandit is in fact the best. Over multiple simulations, this radically increases the variance in performance and thus dampens overall average reward (**Fig. 6c**). However, it is noteworthy that this calculation is made under the assumption that all trials are made with equivalent response times. In the wild, incorrect choices may be less pernicious if they are made rapidly, if biological agents ultimately seek to optimise their reward per unit time (or reward rate).

In our final analysis, we relaxed this assumption and asked how the confirmatory bias affected overall reward *rates*, under the assumption that decisions are drawn to a close after a bounded accumulation process that is described by the drift-diffusion model. This allows us to model not only the choice probabilities but also reaction times.

### Methods of simulations

We simulated a *reinforcement learning drift diffusion model* (RLDDM) in which the drift rate was proportional to the difference in value estimates between the two bandits (Pedersen, Frank, & Biele, 2017), which in turn depends on the update policy (confirmatory, disconfirmatory, or neutral). At each trial, the relative evidence *x* in favour of one of the two options is integrated over time, discretized in finite time step ***i***, until it reaches a threshold ***a***, implying the selection of the favoured option such that:

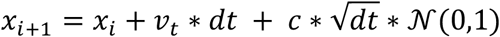

with ***x*_0_**, the initial evidence defined as: 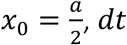 set to 0.001 and ***c*** to 0.1. The drift rate ***v_t_***, is linearly defined from the difference in values such that:

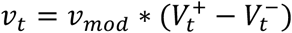

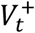 and 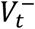 being the values at trial ***t***, of the correct and incorrect options respectively. We used in our simulation a drift rate scaling parameter and a threshold values that make the drift-diffusion model to produce the same choice probabilities as the *softmax* policy with a temperature ***β* = 0.1**. In particular, the probability of making a correct choice by a diffusion model (Bogacz, Brown, Moehlis, Holmes, & Cohen, 2006) is given by:

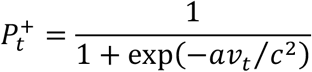

The above probability is equal to that in Equation 5 if ***av_mod_* / *c*^2^ = 1/*β***. Thus, we set 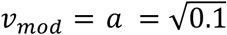. The values are updated exactly the same way as in the *confirmation* model (Equations 2 and 3). We employed the setting with 128 trials, used stable contingencies with reward probabilities equal to ***p*^−^ = 0.35** and ***p*^+^ = 0.65**.

### Results of simulations

When we plotted the overall accuracy of the model, the results closely resemble those from previous analyses, as is to be expected (**Fig. 8a**). When we examined simulated reaction times, we observed that confirmatory learning leads to faster decisions (**Fig. 8b**). This follows naturally from the heightened difference in values estimates for each bandit, as shown in **Fig. 6**. Critically, however, responses were faster for both correct and incorrect trials. This meant that confirmatory biases have the potential to draw decisions to a more rapid close, so that unrewarded errors give way rapidly to new trials which have a chance of yielding reward. This was indeed the case: when we plotted reward rate as a function of confirmatory bias, there was a relative advantage over a neutral bias even for those more extreme confirmatory strategies that were detrimental in terms of accuracy alone (**Fig. 8c**). Thus, even a severe confirmatory bias can be beneficial to reward rates in the setting explored here. However, we note that this may be limited to the case explored here, where the ratio of reward to penalty is greater than one.

**Figure 8.**
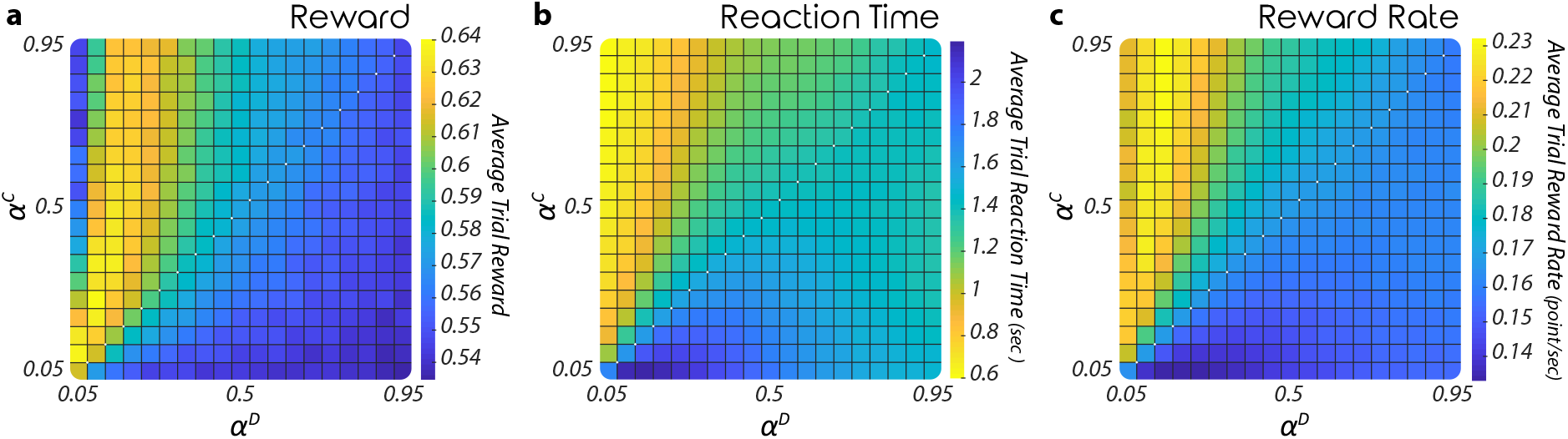
Effect of confirmation bias on reward rate. **a**. The heatmap represents the per trial average reward simulated with the confirmation RLDDM for all learning rates combinations (confirmatory learning rates are represented on the y-axis whereas disconfirmation learning rates are represented on the x-axis). The rewards concern the stable condition with 128 trials and asymmetric contingencies (***p*^−^ = 0.35** and ***p*^+^ = 0.65**) and are averaged across agents. **b**. The heatmap represents the per trial average reaction time estimated with the confirmation RLDDM for all learning rates combinations. **c**. The heatmap represents the per trial average reward rate simulated with the confirmation RLDDM for all learning rates combinations.

## Discussion

Humans have been observed to exhibit confirmatory biases when choosing between stimuli or actions that payout with uncertain probability (Chambon et al., 2019; Palminteri et al., 2017; Schuller et al., 2020). These biases drive participants to update positive outcomes (or those that are better than expected) for chosen options more sharply than negative outcomes, but to reverse this update pattern for the unchosen option. Here, we show through simulations that in an extended range of settings traditionally used in human experiments, this asymmetric update is advantageous in the presence of noise in the decision process. Indeed, agents who exhibited a confirmatory bias, rather than a neutral or disconfirmatory bias, were in most circumstances tested those agents that reaped the largest quantities of reward. This counterintuitive result directly stems from the update process itself that biases the value of the chosen and unchosen options (corresponding overall to the best and worst options respectively), increasing mechanistically their relative distance from each other and ultimately the probability of selecting the best option in the upcoming trials.

Exploring the evolution of action values under confirmatory updates offers an insight into why this occurs. Confirmatory updating has the effect of rendering subjective action values more extreme than their objective counterparts – in other words, options that are estimated to be good are overvalued, and options estimated to be bad are undervalued (**Fig. 6**). This can have both positive and negative effects. The negative effect is that a confirmatory bias can drive a feedback loop whereby poor or mediocre items that are chosen by chance can be falsely updated in a positive direction, leading to them being chosen more often. The positive effect, however, is that where decisions are themselves intrinsically variable (for example, because they are corrupted by Gaussian noise arising during decision-making or motor planning, modelled here with the softmax temperature parameter) overestimation of value makes decisions more robust to decision noise, because random fluctuations in the value estimate at the time of the decision are less likely to reverse a decision away from the better of the two options. The relative strength of these two effects depends on the level of decision noise: within reasonable noise ranges the latter effect outweighs the former and performance benefits overall.

### Relationship to other studies

The results described here thus join a family of recent-reported phenomena whereby decisions that distort or discard information lead to reward-maximising choices under the assumption that decisions are made with finite computational precision – i.e. that decisions are intrinsically noisy (Summerfield & Tsetsos, 2015). For example, when averaging features from a multi-element array to make a category judgment, under the assumption that features are equally diagnostic (and that the decision policy is not itself noisy), then normatively, they should be weighted equally in the choice. However, in the presence of “late” noise, encoding models that overestimate the decision value of elements near the category boundary are reward-maximising, for the same reason as the confirmatory bias here: they inflate the value of ambiguous items away from indifference, and render them robust to noise (Li, Herce Castanon, Solomon, Vandormael, & Summerfield, 2017). A similar phenomenon occurs when comparing gambles defined by different monetary values: utility functions that inflate small values away from indifference (rendering the subjective difference between $2 and $4 greater than the subjective difference between $102 and $104) have a protective effect against decision noise, providing a normative justification for convex utility functions (Juechems, Spitzer, Balaguer, & Summerfield, 2020). Related results have been described in problems that involve sequential sampling in time, where they may account for violations of axiomatic rationality, such as systematically intransitive choices (Tsetsos et al., 2016). Moreover, a bias in how evidence is accumulated within a trial has been shown to increase the accuracy of individual decisions, making the decision variable more extreme and thus less likely to be corrupted by noise (Zhang & Bogacz, 2010).

A recent preprint also reports simulations of the confirmation bias model (Tarantola, Folke, Boldt, Perez, & De Martino, 2021). The simulations in that study paralleled an experimental paradigm reported in that paper, and a confirmation model was simulated for parameters (including softmax temperature) corresponding to those estimated for participant of the study. The simulated agents employing confirmation bias obtained higher average reward than unbiased learners, as well as learners described by other models. Our paper suggests the same conclusion employing a complementary approach in which the models have been simulated in variety of conditions and analysed mathematically.

As mentioned earlier, the consistent benefit of confirmation bias for different reward probabilities (**Fig. 5a** and **5b**) contrasts with the differential effects of biases in learning rates found in a previous study (Caze & van der Meer, 2013). That study considered a standard reinforcement learning task in which feedback is provided only for the chosen option. The study showed that if reward probabilities for both options ***p^i^* < 0.5**, then it is beneficial to have a larger learning rate when 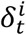 is positive than when it is negative. With such unbalanced learning rates, both ***V*^1^** and ***V*^2^** will be overestimated and critically the difference ***V*^1^ – *V*^2^** will be magnified, so with a noisy choice rule the option with the higher reward probability will be more likely to be selected. By contrast, if both ***p^i^* > 0.5**, then overestimating ***V*^1^** and ***V*^2^** would actually reduce the difference ***V*^1^ – *V*^2^** due to a ceiling effect, because according to Eq 1 reward estimates cannot exceed the maximum reward available, i.e. ***V^i^* ≤ 1**. In this case, it is beneficial to have a larger learning rate when *δ”* is negative than when it positive. In summary, if one assumes that learning rates can differ between rewarded and unrewarded trials, the type of reward-increasing bias depends on magnitude of reward probabilities in a task (Caze & van der Meer, 2013). By contrast, here we show that if one assumes that learning rates can differ between confirmatory and disconfirmatory trials, then confirmation bias tend to increase reward and magnify difference in estimated values both for low reward probabilities (**Fig. 5a** and **7b**) and high reward probabilities (**Fig. 5b** and **7c**).

### Validity of model’s assumptions

Reinforcement learning models fit to human data often assume that choices are stochastic, i.e. that participants fail to choose the most valuable bandit. In standard tasks involving only feedback about the value of the chosen option (factual feedback), some randomness in choices promotes exploration which in turns allows information to be acquired that may be relevant for future decisions. However, our task involves both factual and counterfactual feedback, and so exploration is not required to learn the value of the two bandits. Nevertheless, in some simulations we modelled choices with a softmax rule, which assumes that decisions are corrupted by Gaussian noise, or an *ε*-greedy policy, which introduces lapses to the choice process with a fixed probability. Implicitly, thus, we are committing to the idea that value-guided decisions may be irreducibly noisy even where exploration is not required (Renart & Machens, 2014). Indeed, others have shown that participants continue to make noisy decisions even where counterfactual feedback is available, even if they have attributed that noise to variability in learning rather than choice (Findling, Skvortsova, Dromnelle, Palminteri, & Wyart, 2019).

Due to our assumptions, the present study has a number of limitations. Firstly, we explored the properties of a confirmatory model that has been previously shown to provide a good fit to human data performing a bandit task with factual and counterfactual feedback. However, we acknowledge that this is not the only possible model that could increase reward by enhancing the difference between represented values of options. In principle, any other models producing choice hysteresis might be able to explain these results (Katahira, 2018; Miller, Shenhav, & Ludvig, 2019; Worthy, Pang, & Byrne, 2013). An analysis of these different models and of their respective resilience to decision noise in different settings is beyond the scope of the current study but would be an interesting target for future research. Secondly, the results described here hold assuming a fixed and equal level of stochasticity (e.g. *softmax* temperature) in agents’ behaviours, irrespective of their bias (i.e. the specific combination of learning rates). Relaxing this assumption, an unbiased agent could perform equally well as a biased agent subject to more decision noise. Thus, the benefit of confirmatory learning is relentlessly linked to the level of noise and one level of confirmation bias cannot be thought as being beneficial overall. Thirdly, the present study does not investigate the impact on the performance of other kinds of internal noise such like an update noise (Findling et al., 2019). The latter, instead of perturbing the policy itself, perturbs at each trial the update process of the options’ value (i.e. predictions errors are blurred with a Gaussian noise), and cannot presumably produce a similar increase in performance, having overall no effect on the average difference between these option values.

### Limits to the benefits of biased beliefs

It is important to point out that confirmation bias is beneficial in many but not all circumstances. Although in almost all presented simulations there exists a combination of biased learning rates giving performance that is higher or as good as the best unbiased learner, the optimal learning rates and hence the amount of bias differ depending on task parameters. At the start of the task, a learner usually is unable to know the details of the task a priori, so needs to adopt a certain default combination of learning rates. One could expect that such default learning rates would be determined by past experience or even be to a certain extent influenced by evolution. However such default set of biased learning rates will lead to detrimental effects on performance in certain tasks. For example, a recent study estimated average learning rates of human participants to be ***α^C^* ≈ 0.15** and ***α^D^* ≈ 0.05** (Tarantola et al., 2021) giving a confirmation bias of ***b* ≈ 3**. Although such strong confirmation bias increases reward in many simulated scenarios when decisions are noisy (e.g. **Fig. 3e-h**), it would have a negative effect on performance when decisions are accurately made on the basis of values and in changing environments (e.g. **Fig. 3a-d**). If the default confirmation bias is influenced by evolution, its value is likely to be relatively high, because many of the key decisions of our ancestors had to be quick and thus were noisy due to the speed accuracy trade-off. By contrast, in the modern world, we often can take time to consider important choices, hence the biases that brought evolutionary advantage to our ancestor may not always be beneficial to modern humans.

## Acknowledgements

This work has been supported by MRC grants MC_UU_12024/5, MC_UU_00003/1, BBSRC grant BB/S006338/1, and ERC Consolidator grant 725937.

